# Cortical-subcortical structural connections support transcranial magnetic stimulation engagement of the amygdala

**DOI:** 10.1101/2021.11.12.468411

**Authors:** Valerie J. Sydnor, Matthew Cieslak, Romain Duprat, Joseph Deluisi, Matthew W. Flounders, Hannah Long, Morgan Scully, Nicholas L. Balderston, Yvette I. Sheline, Dani S. Bassett, Theodore D. Satterthwaite, Desmond J. Oathes

## Abstract

The amygdala processes valenced stimuli, influences affective states, and exhibits aberrant activity across anxiety disorders, depression, and PTSD. Interventions that modulate amygdala activity hold promise for treating transdiagnostic affective symptoms. We investigated (*N*=45) whether transcranial magnetic stimulation (TMS) elicits indirect changes in amygdala activity when applied to ventrolateral prefrontal cortex (vlPFC), a region important for affect regulation. Harnessing in-scanner interleaved TMS/functional MRI (fMRI), we reveal that vlPFC neurostimulation evoked acute, dose-dependent modulations of amygdala fMRI BOLD signal. Larger TMS-evoked changes in amygdala fMRI signal were associated with higher fiber density in a vlPFC-amygdala white matter pathway, suggesting this pathway facilitated stimulation-induced communication between cortex and subcortex. This work provides evidence of amygdala engagement by TMS, highlighting stimulation of vlPFC-amygdala circuits as a candidate treatment for affective psychopathology. More broadly, it indicates that targeting cortical-subcortical connections may enhance the impact of TMS on subcortical neural activity and, by extension, subcortex-subserved behaviors.

**Teaser:** Individualized, connectivity-guided transcranial magnetic stimulation modulates the amygdala, demonstrating therapeutic potential.

## Introduction

The amygdala is a critical neural structure for determining an individual’s physiological, emotional, and behavioral responses to affective stimuli. This medial temporal subcortical brain region assigns valence to rewards and threats, facilitates appetitive and aversive conditioning, and influences positive and negative internal affective states as well as associated behaviors (*1*–*4*). Conscious recognition and regulation of amygdala-linked affective states recruits the prefrontal cortex (PFC), including ventrolateral prefrontal (vlPFC) areas subserving voluntary emotional control and affect inhibition (*5*–*11*). Aberrant activity within the amygdala and the vlPFC contributes to symptoms of affective psychopathology observed across many psychiatric diagnoses (*11*–*14*). Indeed, a meta-analysis of task functional MRI data collected from over 11,000 individuals revealed that during emotional processing, patients with mood and anxiety disorders consistently exhibit amygdala hyperactivity and vlPFC hypoactivity—classifying these as two of the most striking and reliable neural phenotypes associated with emotional dysfunction (*11*). Treatments capable of modulating amygdala activity, especially those that simultaneously engage the vlPFC, therefore hold promise for mitigating transdiagnostic affective psychopathology.

Transcranial magnetic stimulation (TMS) is a non-invasive neuromodulation tool that produces changes in neural firing through electromagnetic induction, and that may be capable of eliciting indirect changes in amygdala activity through direct stimulation of functionally or structurally connected cortical locations. Clinically, repetitive TMS administered to the dorsolateral PFC is FDA cleared as a treatment for medication-resistant major depression and obsessive compulsive disorder, and has been studied in clinical trials for post-traumatic stress disorder and anxiety disorders (*15, 16*)—all disorders characterized by amygdala hyperactivity (*11, 13, 14, 17*). Still, despite demonstrated efficacy for many patients with affective symptoms, clinical responses to TMS are variable and not all individuals experience symptom remission. Recent work suggests that the efficacy of prefrontal TMS for affective and post-traumatic stress disorders may depend in part upon the strength of PFC-amygdala functional connections (*18*–*21*), further suggesting that efficacy may vary according to TMS’s ability to alter amygdala functioning. However, to date there is limited direct evidence that prefrontal TMS can specifically modulate amygdala activity (*19, 22, 23*). Furthermore, the extent to which TMS applied to the vlPFC is capable of evoking immediate, reliable changes in amygdala activity remains sparsely investigated, despite the fact that this psychopathology-linked cortical territory is hypothesized to exert top-down control over amygdala neuronal firing (*6, 10*).

TMS alters neural activity by depolarizing somas and large diameter axons, generating action potentials (*24*). Although TMS can only directly depolarize neurons at the cortical surface beneath the device’s magnetic coil (*25, 26*), empirical evidence suggests that TMS can additionally elicit indirect activity changes in “downstream” regions. Perhaps the strongest evidence of this phenomenon comes from motor-evoked potentials: hand muscle electrical potentials recorded in response to TMS of the contralateral motor cortex. These potentials establish that TMS-induced action potentials can propagate along multi-synaptic axonal pathways to elicit activity distant from the cortical site of stimulation (*24*). Additional evidence is provided by studies combining TMS with invasive electrode recordings (*27*) or non-invasive functional MRI (fMRI) recordings (*28*) that have revealed how TMS-induced activity can propagate throughout the brain in a pattern predicted by the stimulated cortex’s structural connectome (*29*).

Combining TMS with fMRI represents a powerful experimental manipulation method, as single pulses of TMS (spTMS) can be delivered inside the MRI scanner interleaved with fMRI functional readouts (spTMS/fMRI). Accordingly, spTMS/fMRI allows one to alter neural activity underneath the TMS coil with stimulation probes while quantitatively measuring effects in the rest of the brain, including in subcortex, constituting a causal “probe-and-measure” approach (*28, 30*). The success of this approach is underpinned by compatibility between TMS-elicited physiological responses and fMRI acquisition properties. Specifically, TMS-elicited changes in neural activity are reliably captured by hemodynamic changes (*25*), which drive the fMRI blood oxygen level-dependent (BOLD) signal. The acute fMRI BOLD response to TMS takes several seconds to peak, thus a time delay can be incorporated prior to the fMRI readout to prevent compromising functional recordings. Moreover, single pulses of TMS briefly evoke neural activity without exerting cumulative effects on firing (*30*), enabling the averaging of single trial fMRI responses to TMS.

In a recent pilot study, our group employed spTMS/fMRI while stimulating a spatially diverse range of lateral PFC sites, and demonstrated feasibility for TMS to evoke downstream changes in the fMRI BOLD signal in the subgenual anterior cingulate cortex and the amygdala (*23*). Critically, in this pilot we observed that stimulation of sites located within or near the vlPFC produced the largest decreases in amygdala BOLD signal. Rhesus macaque tract-tracing work has shown that while the medial PFC is extensively connected to the amygdala (*31, 32*), the majority of lateral PFC areas are only lightly connected—with the exception of the vlPFC (*7*). The vlPFC sends dense, monosynaptic inputs to the amygdala, and thus is the only PFC region with a substantial (as opposed to sparse) amygdala projection that is directly accessible to TMS (*7, 10*). These data support the hypothesis that vlPFC TMS may be particularly capable of modulating amygdala activity due to stimulation-induced action potential propagation along vlPFC-to-amygdala white matter connections. Yet, vlPFC-amygdala structural connections have been scarcely studied in humans (*33*). It therefore remains unknown whether they could comprise one key pathway for cortical-amygdala signal propagation during neuromodulation.

The current study endeavored to causally interrogate whether TMS can exert neuromodulatory effects on the amygdala through the engagement of cortical-subcortical circuits. To accomplish this, we first employed a stimulation-based probe-and-measure approach to validate our preliminary finding that stimulation applied near the vlPFC (“probe”) elicits an acute functional response in the amygdala (“measure”). We next sought to elucidate the structural scaffolding that could allow cortical stimulation to generate a targeted downstream amygdala response. We expected to identify a vlPFC-to-amygdala white matter pathway that is homologous between human and non-human primates; moreover, we hypothesized that pathway properties influencing signal conduction would impact the degree to which TMS affected amygdala activity. The results of our evaluation can be harnessed to guide future TMS protocols that aim to modulate cortical-subcortical circuits involved in affective psychopathology, and are thus readily translatable to TMS clinical trials.

## Results

We leveraged a unique, multimodal dataset to causally probe amygdala fMRI responses to cortical stimulation, and to retrospectively investigate whether the magnitude of response was associated with structural properties of cortical-amygdala white matter connections (**Fig. 1**). This dataset consisted of resting fMRI, structural and diffusion MRI, and in-scanner interleaved spTMS/fMRI data collected from 45 healthy individuals ages 18-55 years (mean age 28 ± 8.6 years; 27 female). This sample of participants was non-overlapping with our pilot TMS/fMRI sample (*23*). To study how non-invasive cortical stimulation affects the amygdala, we applied pulses of TMS in the scanner to individual-specific stimulation sites informed by functional connectivity, and examined fMRI readouts in the subcortex. To explore links between amygdala TMS/fMRI responses and cortical-subcortical structural connectivity, we reconstructed white matter connections between the area of stimulation and the amygdala using fiber orientation distribution (FOD) tractography.

**Fig. 1.**
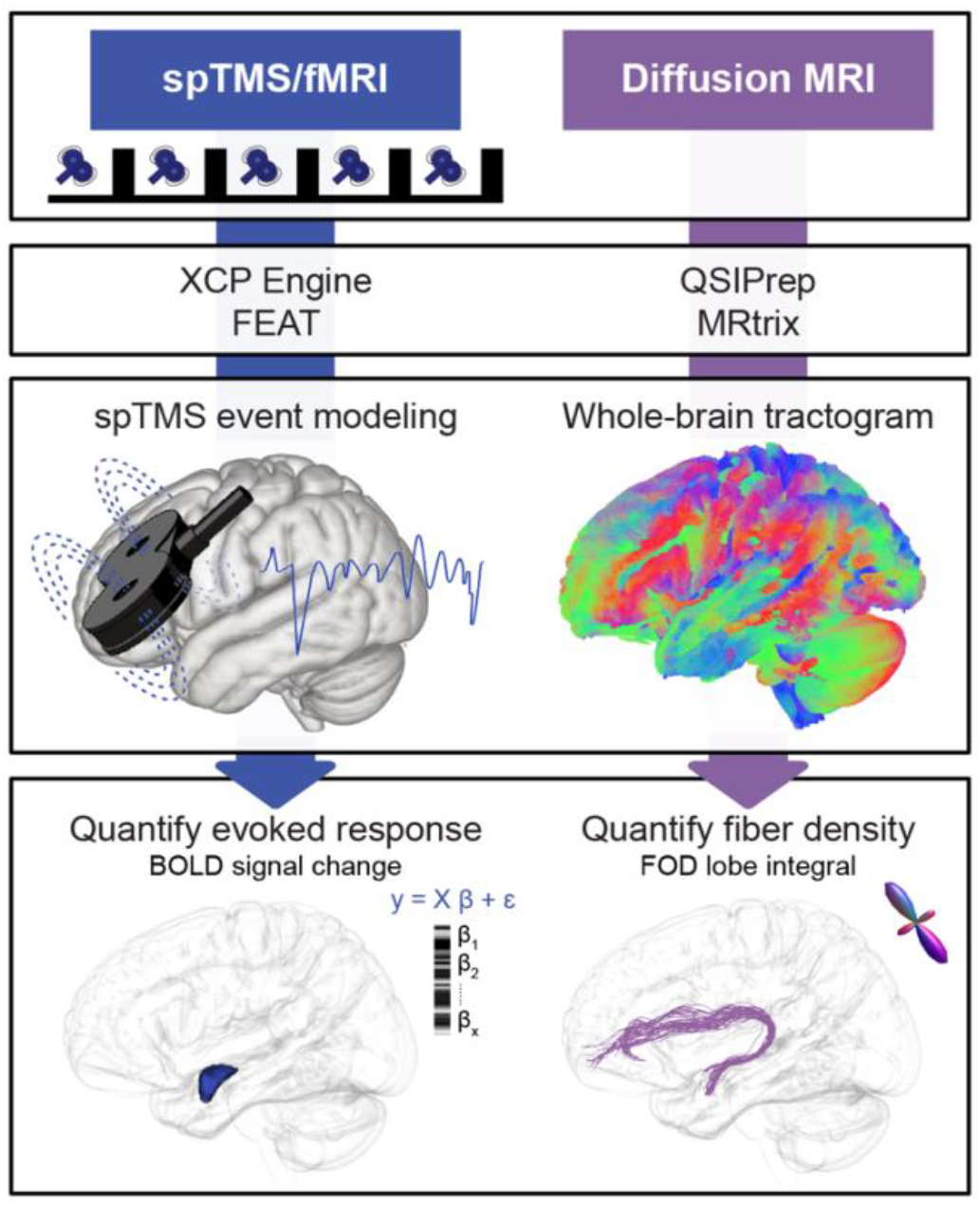
Multimodal Analysis Workflows. *spTMS/fMRI:* Single pulses (sp) of TMS were administered in between fMRI volume acquisitions. TMS pulses were delivered to fMRI-guided, personalized left prefrontal sites of stimulation. Functional timeseries were analyzed with FEAT via XCP Engine’s task module; each TMS pulse was modeled as an instantaneous event. TMS evoked responses were quantified in the left amygdala for each participant by averaging event-related BOLD signal changes induced by stimulation. *Diffusion MRI:* Diffusion data were preprocessed with QSIPrep. Preprocessed images were reconstructed with MRtrix’s single-shell 3-tissue constrained spherical deconvolution pipeline to generate fiber orientation distribution (FOD) images, and a whole-brain tractogram was generated with FOD tractography. A structural pathway connecting the left amygdala to the prefrontal area of TMS stimulation was isolated, and pathway fiber density was quantified.

### Ventrolateral prefrontal cortex TMS modulates fMRI BOLD activity in the amygdala

We employed in-scanner interleaved spTMS/fMRI in order to replicate our prior preliminary study (*23*) in a larger, independent sample and confirm that cortical stimulation exerts neuromodulatory effects on the amygdala, our downstream target of interest. For each participant, a personalized left prefrontal TMS site of stimulation was chosen that exhibited strong functional connectivity to the left amygdala (based on resting fMRI; see Methods) and that was located within, or in closest proximity to, the vlPFC (**Fig. 2A**). A functional connectivity-guided approach was chosen given prior evidence that cortical TMS will elicit larger biobehavioral changes associated with a downstream region, if that region is strongly functionally connected to the cortical stimulation site (*23, 34*–*38*). High functional connectivity sites near the vlPFC were given priority based on our pilot study (*23*), the accessibility of this cortical area to TMS, and monkey tract-tracing work (*7*).

**Fig. 2.**
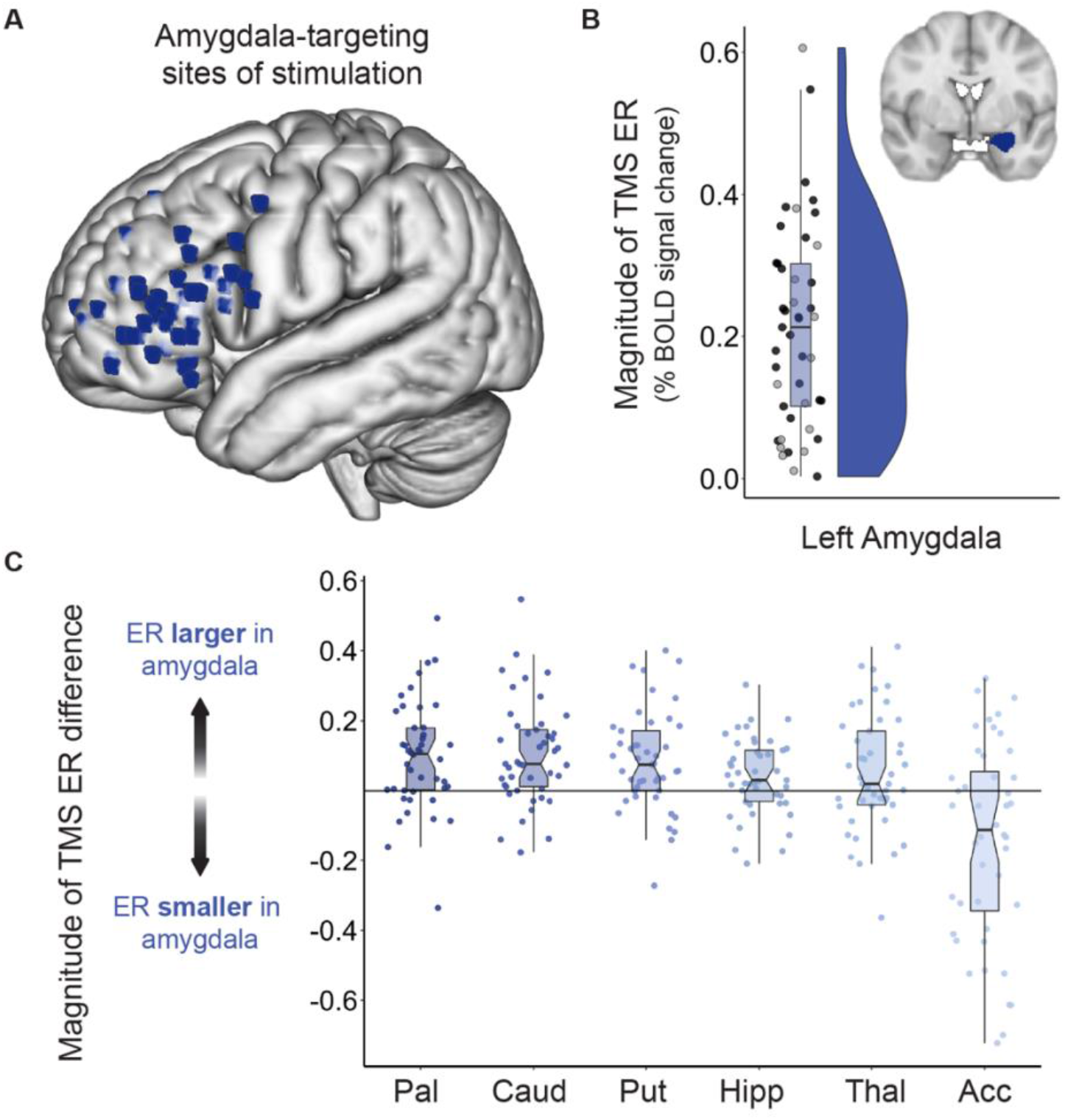
Amygdala BOLD Signal Change Following TMS Administered to vlPFC Connectivity Peaks. **(A)** Each participant’s amygdala-targeting TMS stimulation site visualized in standard (MNI) space. Individual-specific stimulation sites were localized to a left PFC area that was strongly functionally connected to the left amygdala and that was located within the vlPFC (or in closest proximity to the vlPFC of all connectivity peaks). **(B)** TMS elicited a sizable fMRI response in the ipsilateral amygdala. The absolute magnitude of the left amygdala TMS evoked response (TMS ER) is plotted for all participants, along with corresponding box and violin plots. Black circles represent participants (N = 30) that exhibited a negative TMS ER, defined as a TMS-induced decrease in fMRI activity. Grey circles represent participants (N = 15) that exhibited a positive TMS ER, defined as an increase in BOLD signal following stimulation pulses. The box plot displays the median (0.21) and first (0.10) and third (0.30) quantiles of amygdala TMS ERs, with whiskers extending 1.5x the interquartile range. **(C)** TMS evoked responses were overwhelmingly larger in the left amygdala than in other left hemisphere subcortical structures. For each participant, differences in the magnitude (absolute value) of the TMS ER in the left amygdala versus in other left hemisphere subcortical structures were calculated by subtracting each structure’s ER from the amygdala ER; this was done for the left pallidum (Pal), caudate (Caud), putamen (Put), hippocampus (Hipp), thalamus (Thal), and nucleus accumbens (Acc). The magnitude of this evoked response difference is plotted for each subcortical region. Individual participant data points and a notched group boxplot are shown. Data points falling above the y = 0 line indicate that a participant had a larger TMS ER in the amygdala than in the indicated subcortical region.

To empirically assess the impact of single pulses of TMS on ipsilateral amygdala activity, we measured the percent change in BOLD signal elicited by stimulation events, relative to an implicit baseline of no stimulation. We refer to this TMS-evoked change in the fMRI BOLD signal as the TMS “evoked response”. Importantly, both positive and negative evoked responses provide evidence of a transient change in subcortical activity in response to cortical stimulation, and therefore evidence for a cortical-subcortical pathway supporting TMS signal propagation. We thus analyze the unsigned magnitude of the TMS evoked response unless otherwise indicated. Across the 45 study participants, the average absolute value left amygdala evoked response was 0.21% ± 0.14. A BOLD signal change of 0.20% is comparable in magnitude to BOLD effects produced by tasks that functionally engage the amygdala (*39*–*41*), supporting that single pulses of TMS to cortically-accessible sites elicited a functional response in the amygdala (**Fig. 2B**). Examining the direction of each participant’s TMS evoked response revealed that TMS decreased BOLD signal in the amygdala in 30 of 45 individuals, possibly indicative of amygdala inhibition; as a result, the population estimated raw TMS evoked response was negative and significantly different from 0 (*average raw evoked response* = -0.09% ± 0.24, *t*_44_ = -2.51, 95% *CI* = [-0.16 to - 0.02], *p* = 0.0160). Importantly, left amygdala TMS evoked response estimates were highly similar when the amygdala was defined with the Harvard Oxford subcortical atlas (primary approach, reported above) and with individual Freesurfer segmentations, indicating that parcellation choice did not impact quantification of our outcome measure of interest (correlation between approaches: *Pearson’s r* = 0.96, *CI* = [0.93 to 0.98], *p* < .0001).

For all participants, TMS was applied to the left PFC at 120% of the individual’s pre-scan resting motor threshold. However, the distance between the scalp and the cortex—which influences the effective magnitude of cortical stimulation—typically differs between an individual’s primary motor cortex and PFC. Consequently, the strength of neurostimulation ultimately delivered to the PFC may be less than 120% of motor threshold (if scalp-to-cortex distance is greater at the PFC) or greater than 120% (if scalp-to-cortex distance is greater at M1). We therefore corrected the estimated TMS dose for within-individual differences in scalp-to-cortex distance at the stimulation site relative to M1^46^. We observed that the effective strength of neurostimulation varied across participants (distance-adjusted average dose = 110% of motor threshold ± 15%). Moreover, the effective strength of neurostimulation was significantly positively correlated with the magnitude of the left amygdala TMS evoked response (*r*_s_ = 0.35, 95% *CI* = [0.06 to 0.59], *p* = 0.0173), providing evidence for a dose-dependent effect of TMS on amygdala fMRI responses. Absolute stimulator output (% of max) was not correlated with the amygdala evoked response (*r*_*s*_ = -0.09, 95% *CI* = [-0.38 to 0.22], *p* = 0.5764) suggesting that individually-determined motor thresholds corrected for distance provide a more suitable approximation of dose than raw stimulator output.

Next, we sought to assess the specificity of downstream TMS effects within the subcortex. We expected TMS to elicit larger functional responses in the left amygdala than in non-targeted left hemisphere subcortical structures. We thus compared the magnitude of the TMS evoked response in the left amygdala to the magnitude of response in the left caudate, hippocampus, nucleus accumbens, pallidum, putamen, and thalamus (all other Harvard Oxford left hemisphere subcortical structures). Analyses were conducted on absolute valued TMS evoked responses using a within-subjects design, and focused on subcortical regions ipsilateral to the TMS stimulation. We analyzed absolute valued evoked responses as we were interested in whether the overall size of the TMS effect differed between the amygdala and other subcortical structures, rather than whether response direction (positive versus negative) differed between structures. Single pulses of TMS delivered to amygdala functional connectivity peaks within the left vlPFC induced larger changes in BOLD signal in the left amygdala than in the left caudate (*t*_44_ = 4.9, *Cohen’s d* = 0.72, 95% *CI* = [0.06 to 0.15], *p*_FDR_ < 0.0001), the left hippocampus (*t*_44_ = 2.5, *Cohen’s d* = 0.37, 95% *CI* = [0.01 to 0.07], *p*_FDR_ = 0.0201), the left pallidum (*t*_44_ = 4.3, *Cohen’s d* = 0.64, 95% *CI* = [0.05 to 0.14], *p*_FDR_ = 0.0004), the left putamen (*t*_44_ = 4.1, *Cohen’s d* = 0.61, 95% *CI* = [0.04 to 0.13], *p*_FDR_ = 0.0004), and the left thalamus (*t*_44_ = 2.1, *Cohen’s d* = 0.32, 95% *CI* = [0.003 to 0.10], *p*_FDR_= 0.0381) (**Fig. 2C**). In contrast, evoked responses were smaller in magnitude in the left amygdala than in the left nucleus accumbens, suggesting that the amygdala and accumbens may share TMS-targetable cortical representations (*t*_44_ = -3.5, *Cohen’s d* = 0.52, 95% *CI* = [-0.27 to -0.07], *p*_FDR_ = 0.0018, negative accumbens evoked response in 28/45 individuals). To additionally explore whether other subcortical responses to TMS were functionally linked to the amygdala evoked response, we correlated the magnitude of BOLD signal change in the left amygdala with the magnitude of signal change in the aforementioned subcortical structures. Evoked response magnitude in the left amygdala strongly correlated with evoked response magnitude in the left hippocampus (*r*_s_ = 0.59, 95% *CI* = [0.35 to 0.76], *p*_FDR_ = 0.0001), potentially a result of well-known inter-regional connections or spatially proximal cortical inputs. Left amygdala evoked responses did not, however, correlate with evoked responses in the left caudate, nucleus accumbens, pallidum, putamen, or thalamus (all *p*_FDR_ > 0.15), indicating that individual subcortical regions largely display unique functional responses to vlPFC TMS. Together, these findings reveal that the effects of spTMS on the fMRI signal were not only differentiable across subcortical regions, but additionally were almost universally larger in the amygdala—the subcortical structure we aimed to target through cortical functional connectivity.

### A white matter connection provides a pathway for amygdala modulation

We hypothesized that TMS-induced activation of cortical neurons could exert a downstream influence on the amygdala as a result of action potential propagation along a left prefrontal-amygdala white matter pathway. To retrospectively explore this hypothesis, we first created a group TMS stimulation sites mask that combined the 45 individualized amygdala-targeting sites from all participants. We next generated a whole-brain tractogram from a study-specific FOD template, and extracted streamlines with endpoints in the group stimulation mask and the left amygdala. The use of a study-specific FOD template for white matter delineation and feature analysis offers numerous advantages within the context of this study (see Methods for extended discussion). Briefly, compared to individual FOD images, the FOD template has increased signal-to-noise and reduced reconstruction uncertainty, and thereby enables superior tractography algorithm performance and more accurate pathway identification. The template furthermore optimizes anatomical correspondence of the studied pathway across participants, eliminating variability in pathway definitions that can be aliased as between-individual differences in microstructural measures. Finally, the template approach allows for identification of a population representative pathway that can be compared across species.

Our diffusion MRI analysis identified a white matter pathway connecting anterior portions of the left vlPFC to the left amygdala (**Fig. 3**). The human vlPFC-amygdala pathway exhibited close correspondence to the main lateral prefrontal-amygdala pathway identified with invasive tract-tracing in rhesus macaques (*7*). Specifically, non-human primate tract tracing work has shown that the strongest direct (monosynaptic) projection from the lateral PFC to the amygdala originates within area L12 of the vlPFC in macaques, largely corresponding to Brodmann area (BA) 47 and anterior BA 45 in humans (*10*). Using a Brodmann atlas reconstructed in MRI space (*42*), we determined that 60% of pathway streamline endpoints localized to BA47 and BA45 (27% localizing to BA10, 13% to anterior/ventral BA46), confirming that our *in vivo* work recapitulated the spatial pattern of connectivity observed with tract tracing in macaques. Critically, this left vlPFC-amygdala pathway could function as a causal pathway through which TMS-induced modulation of vlPFC activity produced downstream changes in the amygdala.

**Fig. 3:**
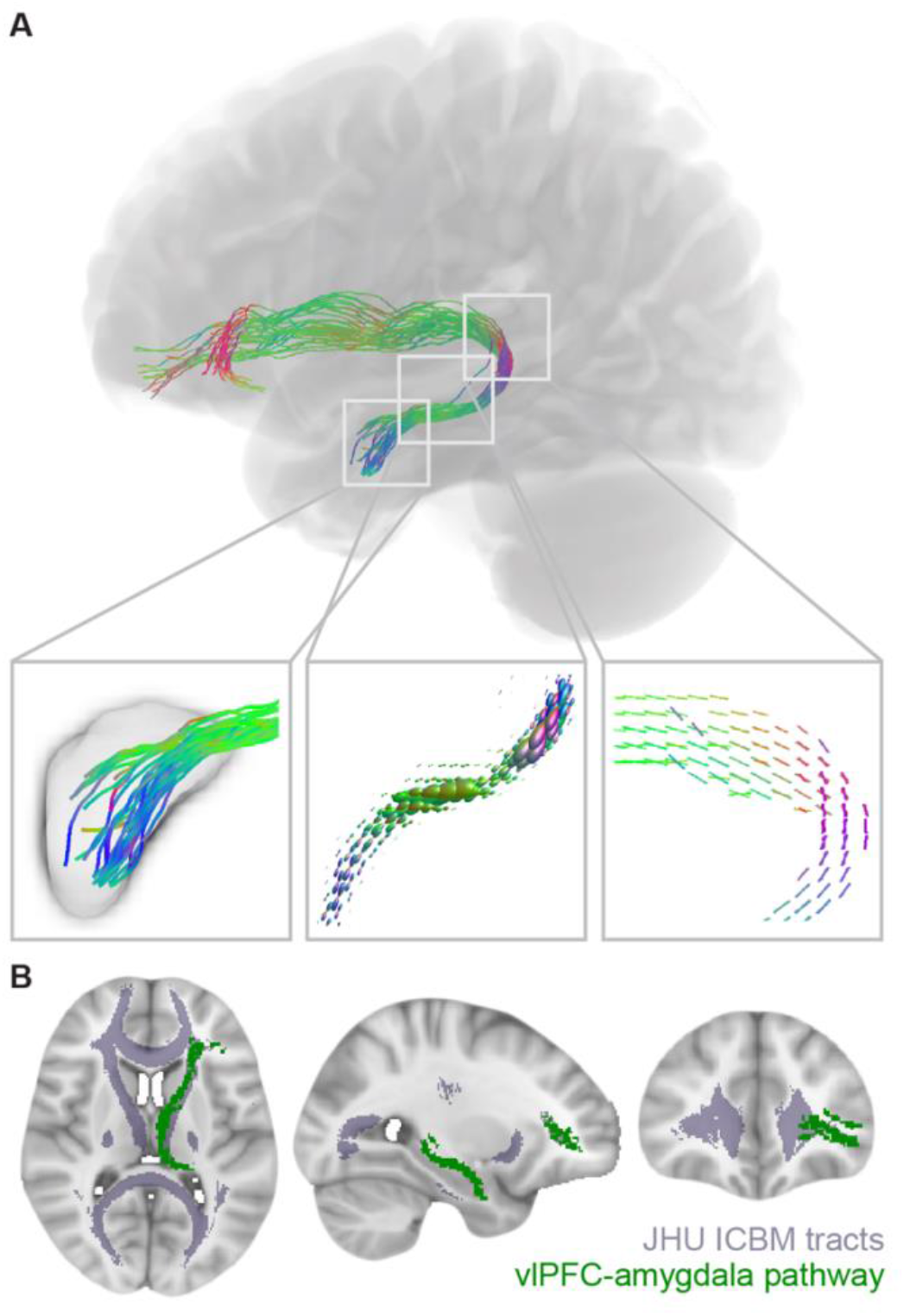
vlPFC-Amygdala White Matter Pathway Anatomy. **(A)** A white matter pathway connecting the left vlPFC stimulation area to the left amygdala could provide a structural scaffold for downstream modulation of the amygdala. This pathway was identified from fiber orientation distribution (FOD) tractography, and pathway streamlines were mapped to individual fiber bundle elements (fixels) for the calculation of fiber density. The left box displays pathway streamlines terminating in the amygdala. The center box displays pathway FODs scaled by fiber density. The right box displays pathway fixels. Colors represent fiber direction. **(B)** The vlPFC-amygdala white matter pathway trajectory is shown. The identified vlPFC-amygdala pathway is shown in green, overlaid on major white matter tracts from the JHU ICBM tract atlas, displayed in purple. The core of the pathway travels with the left anterior thalamic radiation.

### Pathway fiber density is associated with the magnitude of the TMS-evoked amygdala response

If neurostimulation at the cortex leads to downstream changes in the amygdala fMRI signal by engaging this vlPFC-amygdala white matter pathway, then pathway-derived measures should be associated with the amplitude of the amygdala evoked response. In particular, higher pathway fiber density should enable a larger amygdala evoked response by allowing for more effective signal propagation and enhanced cortical input to the amygdala. To quantify fiber density in the vlPFC-amygdala pathway for each study participant, pathway streamlines were mapped to individual fiber bundle elements (also known as “fixels”) in each voxel the pathway traversed, and mean fiber density was estimated across pathway fixels. In support of a circuit-based model of cortical-subcortical TMS signal propagation, individuals with higher fiber density in the left vlPFC-left amygdala white matter pathway exhibited left amygdala TMS evoked responses of significantly greater magnitude (Spearman’s partial correlation, controlling for age: *r*_s.partial_ = 0.36, 95% *CI*= [0.07 to 0.60], *p* = 0.0164) (**Fig. 4A**). Fiber cross-section, a macroscopic, morphological measure of pathway cross-sectional diameter, was not associated with the magnitude of amygdala evoked response (Spearman’s partial correlation, controlling for age and intracranial volume: *r*_s.partial_ = -0.12, 95% *CI* = [-0.40 to 0.19], *p* = 0.4610).

**Fig. 4:**
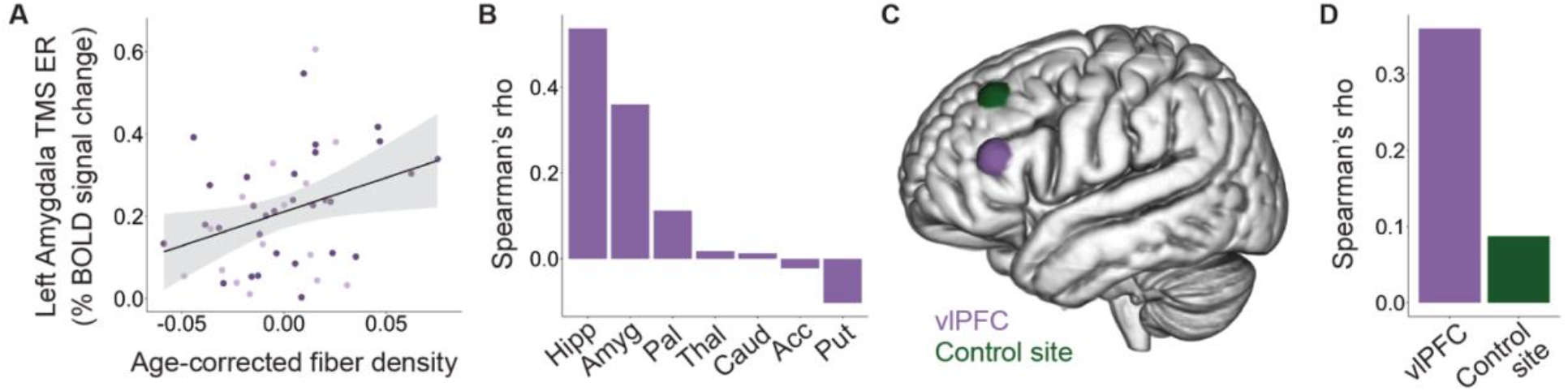
vlPFC-Amygdala White Matter Pathway Fiber Density Impacts Subcortical TMS Evoked Responses. **(A)** Across all participants, higher vlPFC-amygdala white matter pathway fiber density was associated with a greater magnitude left amygdala TMS evoked response (TMS ER). Dark purple circles represent participants that exhibited a negative TMS ER; lighter purple circles represent those that exhibited a positive TMS ER. **(B)** vlPFC-amygdala pathway fiber density was most strongly associated with TMS/fMRI responses in medial temporal subcortical structures, as revealed by the Spearman’s rank partial correlation coefficient (Rho) for each subcortical region. Subcortical regions include the left hippocampus (Hipp), amygdala (Amyg), pallidum (Pal), thalamus (Thal), caudate (Caud), nucleus accumbens (Acc), and putamen (Put). **(C)** In addition to the primary vlPFC spTMS/fMRI scan, each participant received an additional spTMS/fMRI scan during which TMS pulses were applied to an active control site. The intensity-weighted center of gravity of all personalized stimulation sites is shown for vlPFC sites (purple) and control sites (green). **(D)** The strength of the association (Rho) between vlPFC-amygdala pathway fiber density and left amygdala TMS ER magnitude was smaller when TMS was applied to the control site.

In a series of sensitivity analyses, we confirmed that the association between larger left amygdala TMS evoked response and greater left vlPFC-amygdala pathway fiber density was not driven by the strength of neurostimulation, the strength of baseline stimulation site-amygdala functional connectivity, head motion during scanning, head size, or sex. Sensitivity analyses were conducted with independent Spearman’s rank partial correlations controlling for age plus each potential confounder. The association between pathway fiber density and magnitude of the left amygdala TMS evoked response remained significant when controlling for TMS dose (*r*_s.partial_ = 0.31, 95% *CI* = [0.004 to 0.56], *p* = 0.0461) and the TMS site of stimulation in MNI Y and Z coordinates (*r*_s.partial_ = 0.39, 95% *CI* = [0.10 to 0.62], *p* = 0.0108). These observations support that individual-tailored elements of the TMS administration did not explain our finding. Given that stimulation sites were selected based on their resting-state functional connectivity with the left amygdala, we verified that the fiber density-evoked response association could not be attributed to inter-individual differences in the strength of this functional connection (*r*_s.partial_ = 0.31, 95% *CI* = [0.01 to 0.56], *p* = 0.0398). In addition, we showed that the fiber density-evoked response association was not affected by controlling for head motion during the diffusion scan (*r*_s.partial_ = 0.36, 95% *CI* = [0.06 to 0.60], *p* = 0.0179), head motion during the TMS/fMRI scan (*r*_s.partial_ = 0.37, 95% *CI* = [0.08 to 0.61], *p* = 0.0139), total intracranial volume (*r*_s.partial_ = 0.37, 95% *CI* = [0.08 to 0.61], *p* = 0.0142), or participant sex (*r*_s.partial_ = 0.37, 95% *CI* = [0.08 to 0.61], *p* = 0.0140). Finally, we verified that using an alternate method for amygdala parcellation did not have an effect on our findings: the fiber density-evoked response association was significant when the amygdala was identified using participant Freesurfer segmentations (*r*_s.partial_ = 0.36, 95% *CI* = [0.06 to 0.60], *p* = 0.0171), with an effect size equal to that obtained with the Harvard Oxford atlas.

### The identified pathway is differentially associated with neurostimulation-induced subcortical responses

Having demonstrated that the size of the amygdala TMS evoked response was related to fiber density in the delineated pathway, we aimed to establish the specificity of this relationship. We thus examined the association between left vlPFC-amygdala pathway fiber density and spTMS/fMRI BOLD responses in other subcortical structures. Higher vlPFC-amygdala pathway fiber density was also significantly associated with a greater magnitude evoked response in the left hippocampus (Spearman’s partial correlation, controlling for age: *r*_s.partial_ = 0.54, 95% *CI* = [0.28 to 0.72], *p*_FDR_ = 0.0010), in line with the observation that amygdalar and hippocampal TMS evoked responses were correlated. However, vlPFC-amygdala pathway fiber density was not associated with the magnitude of the evoked response in the left caudate, nucleus accumbens, pallidum, putamen, or thalamus (all *p*_FDR_ > 0.90), suggesting substantial specificity for the influence of the pathway on neurostimulation-induced evoked brain responses in the subcortex (**Fig. 4B**).

### Pathway fiber density is not related to the TMS-evoked amygdala response when stimulating a distant control site

In a last analysis, we investigated whether fiber density in the left vlPFC-amygdala pathway was associated with left amygdala TMS evoked response magnitude when TMS was applied to a spatially distant, active control site not thought to have direct connections to the amygdala. Control site spTMS/fMRI data were acquired from all individuals on the same day as the amygdala-targeting spTMS/fMRI data, in a pseudorandom counter-balanced design. Control sites of stimulation were located dorsal and posterior to the amygdala-targeting stimulation sites; control and amygdala-targeting sites were located on average 4.4 (± 1.5) cm apart (**Fig. 4C**). Single pulses of TMS applied to the control site elicited an average absolute value left amygdala evoked response of 0.19% ± 0.25, with a negative evoked response observed in 28 of 45 participants. The absolute magnitude of the left amygdala evoked response was larger when stimulating the vlPFC than when stimulating the control site in 62% of participants (0.15% larger on average), although this did not represent a statistically significant difference in magnitude (*V* = 653, 95% *CI* = [-0.01 to 0.10], *p* = 0.1284). As expected, we did not identify structural connections between the amygdala and control TMS sites (using a group mask that combined all participants’ control stimulation sites), suggesting that control site TMS could have affected amygdala activity by engaging poly-synaptic connections (*10*). Finally, we hypothesized that because control site stimulation would be unlikely to directly engage the left vlPFC-amygdala pathway, there would not be a relationship between the microstructure of this white matter pathway and changes in left amygdala activity elicited by control site TMS. Indeed, when TMS was applied to the control site, vlPFC-amygdala pathway fiber density was not significantly associated with the magnitude of the left amygdala TMS evoked response (Spearman’s partial correlation, controlling for age: *r*_s.partial_ = 0.09, 95% *CI* = [-0.22 to 0.38], *p* = 0.5729) (**Fig. 4D**).

## Discussion

A substantial percentage of individuals experiencing affective psychiatric symptoms do not experience a satisfactory clinical response to currently available treatments, necessitating modified or new treatment protocols. A promising, experimental therapeutics based approach for developing translatable protocols is to identify interventions that are capable of engaging brain regions (targets) strongly linked to symptomatology (*43*), such as the amygdala. TMS represents both a psychiatric treatment that can be further optimized and—when combined with fMRI—a tool for measuring target engagement. In the present study, we harnessed interleaved spTMS/fMRI to examine the impact of prefrontal TMS on the amygdala, and established that single pulses of TMS delivered within or near the vlPFC elicit acute, dose-dependent modulations of the amygdala fMRI BOLD signal. We additionally delineated a phylogenetically-conserved white matter pathway connecting the vlPFC to the amygdala with the potential to transmit TMS-induced neural activity from the stimulated cortical surface to the medial temporal lobe. Higher fiber density in the identified pathway was associated with larger magnitude TMS-evoked fMRI BOLD responses in the amygdala when stimulating the vlPFC, but not when stimulating an active control site, supporting a specific role for this pathway in vlPFC-to-amygdala TMS signal transduction. Broadly, this spTMS/fMRI probe-and-measure study demonstrates proof of amygdala engagement by TMS, and furthermore highlights a potential structural mechanism facilitating engagement of this subcortical target.

Studies investigating the neural bases of psychiatric treatment response have repeatedly reported that reductions in depressive, anxiety, obsessive-compulsive, and post-traumatic stress symptoms occur concomitantly with a normalization of amygdala activity (*17, 40, 44*–*48*) Associations between clinical improvement and modified amygdala functioning have been observed following treatment with psychotropics, cognitive behavioral therapy, electroconvulsive therapy, and surgical interventions, convergently suggesting that neuromodulation of the amygdala may facilitate efficacious reductions in affective psychopathology. Here we provide neuromodulation-relevant evidence that TMS applied to left prefrontal-amygdala functional connectivity peaks can evoke a downstream change in ipsilateral amygdala fMRI activity, with a degree of anatomical specificity. In particular, our data show that TMS tended to induce a negative evoked response, or a decrease in BOLD signal, in the amygdala. Given that heightened amygdala BOLD activity is consistently observed in persons with psychiatric disorders (*11, 13, 14*) this may putatively be the clinically preferred direction of TMS response in this region. It is possible, however, that enhancing amygdala activity may prove beneficial in some contexts. Increases in amygdala neuronal activity are required, for example, for the extinction of conditioned fear (*4, 49, 50*). Accordingly, it will be important for future work to examine whether positive versus negative amygdala TMS evoked responses are associated with differential behavioral or clinical outcomes, for example with dissociable changes in fear conditioning, negative affect, valence evaluation, or emotion regulation.

This study additionally demonstrated that non-invasive brain stimulation engages the amygdala when specifically applied to the vlPFC, a cortical region that is recruited for emotional regulation and transdiagnostically hypoactive in patients with affective psychopathology (*11, 14*). This represents a replication of our prior preliminary study (*23*) and provides further brain-based evidence identifying the vlPFC territory with axonal projections to the amygdala as a candidate TMS treatment target for affective psychiatric disorders. Behavior-based evidence corroborating the potential utility of brain stimulation through this circuit is offered by two independent investigations into vlPFC stimulation. In the first investigation, vlPFC TMS facilitated the regulation and reduction of negative emotions in healthy individuals (*51*). In the second, direct electrode stimulation of the anterior vlPFC produced acute improvements in mood in individuals with depression (*52*). Complementary evidence thus indicates that vlPFC stimulation can impact both neural and clinical features that are disrupted in mood and anxiety disorders. Of note, the medial PFC is also robustly implicated in affective symptomatology and interconnected with the amygdala, and is thus a cortical territory of interest for some forms of stimulation-based treatments in psychiatry (*20, 21, 31, 32, 53, 54*). However the induced electric field produced by TMS cannot directly penetrate the medial PFC, highlighting the practical utility of stimulating the vlPFC with TMS to preferentially engage the amygdala.

The vlPFC’s structural pathway to the amygdala may allow TMS to synchronously affect neural activity in both of these regions due to direct depolarization of their axonal connections. The putative importance of directly modulating this vlPFC-amygdala pathway is informed by reports from deep brain stimulation (DBS) in psychiatry: subcortical DBS is significantly more effective at reducing psychiatric symptoms when the electrodes contact cortical-subcortical white matter connections (*54*–*58*). The relevance of this pathway is further underscored by the finding that higher pathway fiber density was associated with larger TMS-induced fMRI activity modulations—yet only within medial temporal lobe subcortical structures, and only when stimulating the vlPFC. Our diffusion MRI findings thus provide *in vivo* evidence that greater white matter conductance enhances the ability of TMS-elicited neural signals to travel to distant brain regions, with white matter connectivity profiles in part determining the pathway of signal travel. A central role for white matter in shaping downstream responses to TMS highlights the potential for structural connectivity to be harnessed to engage psychopathology-relevant subcortical structures effectively and focally.

To date, cortical-subcortical functional connectivity has principally been used to target subcortical structures with TMS, with a notable degree of clinical success within the context of major depression (*35, 37, 38, 59*). Nevertheless, cortical functional connectivity weights for a given subcortical target can vary over time in the same individual, impacting the reproducibility of TMS stimulation site selection (*60*). White matter pathways form by early childhood and remain in existence for one’s lifetime, thus potentially offering a complementary approach to guide TMS coil positioning. Integrative strategies harnessing both structural and functional connectivity are thus particularly worthy of future study. These personalizable, precision connectomics strategies could be applied not just to enhance the ability of TMS to modulate the amygdala, but to reach additional subcortical targets that contribute to diverse forms of psychopathology.

The present work must be considered within the context of conventional limitations associated with the *in vivo* neuroimaging measures employed. TMS-evoked fMRI BOLD responses only indirectly index changes in neuronal activity, and can additionally be influenced by changes in metabolism, cerebrovascular reactivity, and neurovascular coupling. The white matter fiber density measure employed here is not an explicit measure of the number of axons present. However, increases in axon count or packing density (or, potentially, decreases in extracellular space) within a fixel will be reflected as an increase in fiber density. As with all tractography methods, we cannot unequivocally determine whether the structural pathway identified between the left vlPFC and the left amygdala represents a direct or a polysynaptic connection, although tract-tracing data compellingly suggest it may be monosynaptic (*7*). Two additional limitations represent key avenues for future investigations. First, this study was not designed to identify factors related to whether an individual exhibited a positive or negative TMS evoked response in the amygdala; future work should explore the impact of TMS stimulation parameters, TMS coil orientation, and the participant’s cognitive or emotional state on response directionality (*61*–*64*). Second, we employed a retrospective study design to examine associations between vlPFC-amygdala white matter pathway features and TMS evoked BOLD responses. Consequently, the TMS coil was not always precisely positioned over the center of the pathway’s cortical fiber terminations, as could be accomplished in a future, prospective structural connectivity-based targeting study.

This study demonstrates that spTMS/fMRI and diffusion MRI can be jointly harnessed to examine how cortical neurostimulation affects activity in brain regions associated with the manifestation and treatment of transdiagnostic affective psychopathologies. Our findings underscore the relevance of examining downstream, subcortical effects of TMS, and the importance of mapping causal circuits underlying these effects. Circuit mapping approaches have been applied in DBS to increase the clinical efficacy of stimulation protocols (*54*–*58*), and, as shown here, can be translated to TMS with the goal of informing treatment protocols. Ultimately, integrating insights derived from spTMS/fMRI brain-based readouts and diffusion-based connectivity into TMS protocols may help to increase the impact of TMS on both brain activity and behavior—thus enhancing the efficacy of therapeutic TMS for psychiatric conditions.

## Materials and Methods

### Experimental Design

Healthy participants ages 18 to 55 years with no present or prior reported neurological or psychiatric conditions and no psychotropic medication use participated in this study. All participants gave informed consent prior to study participation, and all procedures were approved by the University of Pennsylvania Institutional Review Board. All research procedures were performed in accordance with the Declaration of Helsinki. The 45 individuals included in the final study sample had T1-weighted, diffusion, resting state fMRI, and interleaved spTMS/fMRI data (both amygdala-targeting site and control site data) that passed stringent visual and quantitative quality control procedures. Nine additional individuals had neuroimaging data acquired at the time of analysis but were excluded from the study due to excessive motion or image artifacts. Exclusion criteria included an average relative motion root mean square > 0.15 during spTMS/fMRI scans (4 excluded) or an average framewise displacement > 0.20 during the diffusion scan coupled with motion-induced patterned slice drop out observed in diffusion gradients (2 excluded) or reconstructed FOD images (3 excluded). All neuroimaging data were acquired on the same 3 Tesla Siemens Prisma MRI scanner over two separate scanning days, including a baseline scan day and a TMS/fMRI scan day. During the baseline scan, data from resting state fMRI, diffusion MRI, and T1-weighted structural MRI sequences were acquired. The resting state data were collected in order to identify participant-specific regions in or near the left vlPFC that exhibited strong functional connectivity to the left amygdala. These personalized PFC-amygdala functional connectivity peaks were used as sites of stimulation on the TMS/fMRI scan day. The diffusion MRI data were utilized to retrospectively evaluate the hypothesis that TMS-induced changes in cortical activity could have a downstream effect on amygdala activity due to a prefrontal-amygdala white matter pathway. Baseline T1-weighted data were used in both fMRI and diffusion analysis streams. During the TMS/fMRI scan day, TMS was applied in the scanner interleaved with fMRI volume acquisitions in order to quantify evoked changes in amygdala activity in response to single pulses of cortical neurostimulation.

### TMS site of stimulation localization: resting state functional MRI

Baseline resting state fMRI data were collected to enable fMRI-guided selection of TMS sites of stimulation. Two baseline eyes-open (fixation cross focus) multiband resting state fMRI scans were acquired with reverse phase encoding directions in 72 interleaved axial slices with the following acquisition parameters: repetition time = 800 ms, echo time = 37 ms, flip angle = 52°, field of view = 208 mm, voxel size = 2 mm^3^, 420 measurements. A multi-echo T1-weighted MPRAGE scan was additionally acquired with the following parameters: repetition time = 2400 ms, echo time = 2.24 ms, inversion time = 1060 ms, flip angle = 8°, voxel size = 0.8 mm^3^, field of view = 256 mm, slices = 208, PAT mode GRAPPA.

T1-weighted scans were processed with the Advanced Normalization Tools (ANTS) Cortical Thickness Pipeline (*65*). Resting state fMRI data were preprocessed with the eXtensible Connectivity Pipeline Engine (XCP Engine) (*66*) in order to implement a well validated, top performing pipeline for mitigating motion-related artifacts and noise in fMRI data (*67*). Preprocessing steps for the fMRI data included merging of AP and PA acquisitions, removal of the first 2 volumes from each run to allow for scanner equilibration, realignment of all volumes to an average reference volume, identification and interpolation of time series intensity outliers with AFNI’s 3dDespike, demeaning and both linear and polynomial detrending, and registration of fMRI data to T1-weighted data using boundary-based registration. Artifactual variance was modeled as a linear combination of 36 parameters, including 6 motion-related realignment parameters estimated during preprocessing, the mean signal in deep white matter, the mean signal in the cerebrospinal fluid compartment, the mean signal across the entire brain, the first temporal derivatives of the prior 9 parameters, and quadratic terms of both the prior 9 parameters and their derivatives. These 36 nuisance parameters were regressed from the BOLD signal with a general linear model. Last, simultaneous with confound regression, the BOLD time series and the artifactual model time series were temporally filtered (first-order Butterworth) using high-pass-only and low-pass-only filters of > 0.01 Hz and < 0.08 Hz, respectively. In order to transform preprocessed fMRI data to MNI space for functional connectivity analysis, T1-weighted images were non-linearly registered to the MNI T1 template using ANTS symmetric diffeomorphic image normalization (SyN), and transforms were applied to the functional image.

Following preprocessing, functional connectivity—defined as the Fisher’s z-transformed Pearson correlation coefficient between two BOLD time series—was computed between left frontal cortex voxels and a left amygdala seed, as in prior work (*23*). The amygdala functional connectivity map was then transformed back to participant T1 space and stereotaxically visualized on each participant’s curvilinear reconstructed brain surface with a state-of-the-art neuronavigation system (Brainsight; Rogue Research, Montreal, Quebec, Canada). This process allowed for identification of a cortically-accessible stimulation site for the in-scanner TMS/fMRI session that exhibited high functional connectivity to the left amygdala and that localized to (or nearest to) the vlPFC. On the TMS/fMRI scan day, the Brainsight neuronavigation system was used to pinpoint the location on the scalp (marked on a secured lycra swim cap) perpendicular to the amygdala-targeting cortical stimulation site; the TMS coil was centered on this location. Preprocessed resting state fMRI data were additionally used to define the active control sites of stimulation for this study. Each participant’s control site was located in the left middle or superior frontal gyrus, distant from the amygdala-targeting site (4.4 cm on average). Control sites were selected for exhibiting high functional connectivity to the left subgenual anterior cingulate cortex, rather than selected for low functional connectivity to the amygdala per se. Control sites were selected in Brainsight using seed-to-voxel functional connectivity maps generated with a subgenual seed, as in prior work (*23*).

### TMS evoked response quantification: in-scanner, interleaved spTMS/functional MRI

We acquired in-scanner interleaved spTMS/fMRI scans while applying TMS to the scalp location that focused stimulation to PFC-amygdala functional connectivity peaks located in closest proximity to the vlPFC. An MRI-compatible TMS coil (Magventure MRI-B91 air cooled coil) was positioned to induce a posterior to anterior current, and stimulation intensity was applied at 120% of an individual’s resting motor threshold. Resting motor threshold was determined within the MRI room immediately prior to scanning, and defined as the stimulation intensity required to elicit visually observable motor activity in the right hand (in abductor pollicis brevis or first dorsal interosseous muscles) on 5 out of 10 consecutive trials. spTMS/fMRI scans were acquired using a TMS-compatible birdcage head coil (RAPID quad T/R single channel; Rimpar, Germany). During scanning, the MRI-B91 TMS coil was connected to a Magpro X100 stimulator (Magventure Farum, Denmark) and held firmly in place by a custom-built TMS coil holder. The spTMS/fMRI acquisition parameters included: repetition time = 2000 ms, echo time = 30 ms, flip angle = 75°, field of view = 192 mm, voxels = 3×3×4 mm, 32 interleaved axial slices, 174 measurements. Transistor-transitor logic (TTL) trigger pulses sent through a parallel port with E-prime 2.0 (Psychology Software Tools, Sharpsburg, Pennsylvania, USA) were used to control the timing of fMRI volume acquisitions and single TMS pulses^34^. Individual fMRI volume acquisitions were spaced by a 400 ms window during which a single pulse of TMS was delivered (triggered at 200 ms). This temporal spacing allows for administration of TMS pulses in a manner that does not contaminate the magnetic field during the subsequent volume acquisition. The TMS/fMRI scan was broken into 12 spTMS/fMRI mini-blocks throughout which a total of 71 TMS pulses were administered. Each mini-block consisted of 7 400-ms windows during which TMS could be delivered interleaved with 7 fMRI volume acquisitions. TMS was administered during 5 to 7 of the mini-block 400 ms windows in order to incorporate 0-2 catch trials, preventing prediction of when TMS would be delivered. Mini-blocks were separated by 7 fMRI volume acquisitions.

Amygdala-targeting spTMS/fMRI data were preprocessed with XCP Engine’s task module, which executes the FMRI Expert Analysis Tool (FEAT, version 6.0.0). The functional data were motion corrected using six standard motion regressors with FSL MCFLIRT, high-pass temporally filtered (cut off of 100), spatially smoothed (5 mm FWHM kernel), registered to baseline T1-weighted images using boundary-based registration, and transformed to MNI space using pre-computed T1-MNI registration transforms. For event modeling, each TMS pulse was considered an instantaneous event and convolved with a gamma-shaped hemodynamic response function. Following model estimation, parameter estimates and contrast values were used to calculate the percent change in BOLD signal from no stimulation (implicit baseline) to stimulation. The average percent BOLD signal change was then quantified in left hemisphere subcortical structures using the Harvard Oxford subcortical atlas, yielding region-specific TMS evoked responses. A positive evoked response indicates a TMS-induced increase in BOLD signal, whereas a negative evoked response indicates a TMS-induced decrease in BOLD signal. The magnitude of the evoked response indexes the overall size of the response regardless of direction (i.e., the absolute value), and provides insight into the strength of the functional response elicited by neurostimulation—thereby capturing the main neurobiological effect of interest in this study. On the TMS/fMRI scan day, a second spTMS/fMRI scan was acquired in a counter-balanced design with TMS targeted to the control site. The control site spTMS/fMRI scan was acquired and processed exactly as detailed above for the amygdala-targeting scan.

### Prefrontal-amygdala white matter pathway delineation: diffusion MRI

Our diffusion MRI analytic workflow sought to determine whether white matter connections originating in the area of cortical stimulation could serve as pathways for TMS-induced signal travel to the amygdala. Diffusion data were acquired in 64 gradient directions with b = 1000 s/mm^2^ (and one b = 0 volume) with the following parameters: repetition time = 4000 ms, echo time = 72.60 ms, flip angle = 90°, voxel size = 2 mm^3^, slice number = 76. The data were preprocessed with QSIPrep 0.6.3RC3, a containerized pipeline that integrates algorithms from diverse software and implements critical preprocessing steps with the best tools available in the field (*68*). In QSIPrep, the data were denoised with Marchenko-Pastur principal component analysis (MP-PCA) (*69*), head motion and eddy currents were corrected using FSL eddy with outlier replacement (*70*), and susceptibility distortions were corrected with fieldmaps generated from magnitude and phase difference images. A non-diffusion weighted reference image (b=0) from the preprocessed diffusion data was registered to a skull-stripped, AC-PC aligned T1-weighted image. A single BSpline interpolation was then applied to both upsample the diffusion data to a 1.3 mm^3^ voxel resolution and align it with the AC-PC realigned T1-weighted image. All subsequent diffusion analyses, including signal reconstruction with a higher-order diffusion model, tractography, and fixel metric quantification, were employed following recommended pipelines in MRtrix3 (*71*) (https://mrtrix.readthedocs.io/en/3.0.0/fixel_based_analysis/st_fibre_density_cross-section.html) using MRtrix3Tissue version 5.2.8 (https://3Tissue.github.io). With MRtrix3Tissue, diffusion images were reconstructed with single-shell 3-tissue constrained spherical deconvolution (*72*) using a set of group-average white matter, gray matter, and cerebrospinal fluid response functions estimated with the *dhollander* algorithm (*73*). Constrained spherical deconvolution was implemented for reconstruction as it allows for the delineation of multiple anatomically-accurate fiber populations per voxel through estimation of a fiber orientation distribution. Each set of antipodally symmetric FOD lobes represents a distinct fiber population; the shape and amplitude of the lobes provides information about fiber microstructure. Critically, the use of 3-tissue response functions during deconvolution removes extra-axonal signal contributions from gray matter and cerebrospinal fluid, increasing the precision of the FOD and the biological specificity of the fiber density metric.

Following construction of participant FOD images, images underwent 3-tissue bias field correction and global intensity normalization to ensure that absolute FOD amplitudes were directly comparable across all images. A study-specific FOD template was then created using normalized data from all participants. The template was used to conduct FOD-based tractography (iFOD2 algorithm, MRtrix3 default parameters, 2.5 million streamlines), producing a whole-brain tractogram (*74*). Subsequently, streamlines with endpoints in a group TMS stimulation sites mask and a left amygdala mask were extracted—delineating a vlPFC-amygdala structural pathway that could support TMS-induced actional potential propagation. The TMS stimulation sites mask was a study-specific mask comprised of dilated amygdala-targeting TMS sites (Fig. 2A). The left amygdala was delineated using the Harvard Oxford subcortical atlas. In order to quantify participant-specific measures within the fiber populations that constitute the extracted vlPFC-amygdala pathway, a fixel-based analysis pipeline was implemented as previously described in detail (*75*). vlPFC-amygdala pathway streamlines were mapped to individual fixels, and each participant’s average fiber density and average fiber cross-section was calculated across fixels corresponding to the pathway. A primary streamline-to-fixel mapping threshold of 5 streamlines was used to ensure the robustness of the pathway, in accordance with prior publications (*76*). We verified, however, that findings were reproducible at mapping thresholds of 2, 4, 6, and 8. Fiber density, quantified by the integral of the FOD lobe, is a microstructural measure of a pathway’s intra-axonal volume per unit volume of tissue (accounting for crossing fibers) that is sensitive to axon count and packing density (*77*). Fiber cross-section is a morphological measure, computed from the Jacobian determinant of a participant-to-template non-linear warp, that is affected by pathway diameter. Fiber cross-section was log transformed to ensure normality, as advised in the MRtrix3 documentation.

Conducting tractography on a study-specific FOD template rather than on individual participant FOD images confers numerous advantages within the framework of the present study. As compared to individual FOD images, the study-specific FOD template has greatly enhanced signal-to-noise and reduced uncertainty associated with each FOD (*77*). The superior FOD reconstruction quality supports improved tractography performance and lowers susceptibility to spurious streamlines, thus likely increasing the anatomical validity of identified pathways. Extracting streamlines of interest based on a study-specific tractogram also ensures that only white matter pathways that are well represented across the entire study population are analyzed. The delineation of tracts that are highly representative of the population allows for both more apposite across-species comparisons (i.e., between human tractography and macaque tract-tracing) and for more appropriate comparisons across individuals. Specifically, by optimizing anatomical correspondence of the vlPFC-amygdala pathway across individuals, the template approach ensures that inter-individual differences in pathway fiber density cannot simply be attributed to differences in delineation of the pathway itself. This is critical as past work from our group has shown how variability in the extraction of a white matter pathway’s streamlines can produce artifactual differences in microstructural measures of interest (*78*). Finally, the template approach additionally enables the examination of macrostructural morphological measures that are based on the participant-to-template FOD warp.

## Statistical Analysis

Statistics were conducted in R 4.0.2. A two-sided, one-sample t-test was conducted to determine if, on average, raw TMS evoked responses in the left amygdala were significantly greater or less than 0 when stimulating near the vlPFC. Differences between amygdala evoked response magnitude and evoked response magnitude in other subcortical structures were evaluated with two-sided, paired-samples t-tests, after confirming normality of paired differences. T-tests were performed with the t.test function (stats package in R); corresponding effect sizes were estimated with the cohensD function (lsr package). To compare left amygdala evoked response magnitudes when targeting vlPFC sites versus active control sites, a two-sided paired Wilcoxon signed rank test (*mu* = 0) was utilized, given that the paired differences were non-normally distributed (wilcox.test function, stats package). Non-parametric Spearman’s rank correlations (denoted by *r*_s_) were carried out to determine how correlated the magnitude of the amygdala TMS evoked response was with TMS dose and with response magnitude in other subcortical structures. Spearman’s rank partial correlations (denoted by *r*_s.partial_) controlling for age were employed to quantify associations between TMS evoked response magnitude and white matter fiber density or fiber cross section. The fiber cross-section analysis additionally included intracranial volume as a covariate, given that this morphological measure is strongly correlated with brain size (*79*). Full and partial Spearman’s correlations were implemented with cor.test (stats package in R) and pcor.test functions (ppcor package), respectively; correlation coefficient confidence intervals were estimated with the cor_to_ci function (correlation package). Throughout, false discovery rate correction was applied to correct for multiple comparisons (denoted by *p*_FDR_) when multiple subcortical structures were examined in an analysis.

## Code Availability

All study analytic code and a guide to code implementation are available at https://pennlinc.github.io/ZAPR01_dMRI_TMSfMRI/.

## Funding

National Science Foundation Graduate Research Fellowship DGE-1845298 (VJS)

National Institutes of Mental Health R01MH111886 (DJO)

National Institutes of Mental Health RF1MH116920 (TDS, DSB, DJO)

## Author contributions

VJS and DJO conceived of the study. RD, JD, HL, and MS acquired the MRI data. RD and MWF processed the structural and resting state functional MRI data and identified TMS stimulation sites, with guidance from DJO. VJS processed the diffusion MRI data, with guidance from MC. VJS implemented all statistical analyses and generated all figures. MC conducted an internal code review and technical replication. NLB, YIS, DSB, TDS, and DJO helped with data interpretation and clinical applicability. VJS and DJO wrote the manuscript. All authors revised the manuscript.

## Data and materials availability

The neuroimaging data collected and analyzed for the current study are available upon reasonable request with a data use agreement. All code written for TMS evoked response quantification, diffusion MRI analysis, statistical analysis, and visualization is available at https://github.com/PennLINC/ZAPR01_dMRI_TMSfMRI.

